# Reproducible Tools and Enhanced Computational Workflows for Batch Effect Evaluation of High-Throughput Data Using BatchQC

**DOI:** 10.64898/2026.02.12.705115

**Authors:** Jessica K Anderson, Jiwei Zhang, Xinshou Ge, Howard Fan, Yaoan Leng, Michael Silverstein, Regan Conrad, Zhaorong Li, Evan Holmes, Solomon S. Joseph, Sean Lu, Russell T Shinohara, Tenglong Li, W. Evan Johnson, Alzheimer’s Disease Neuroimaging Initiative

## Abstract

Batch effect correction is a common and often necessary step in data analysis to reduce bias due to technical and experimental factors when combining multiple batches of data. The severity of the batch effects dictates the correction strategy; therefore, a careful assessment of each dataset’s batch effects is necessary. BatchQC is an R package that provides reproducible tools and visualizations for quantitatively and qualitatively addressing batch effects across a broad range of data types. BatchQC integrates with standardized Bioconductor data structures and features an object-oriented design, enabling the application of workflows that can freely evaluate and process data within and outside the package tools. Common batch evaluation methods, along with novel quantitative metrics, help determine the benefits of batch correction for each dataset and enable direct comparisons between methods. Here, we present BatchQC as the first comprehensive batch-correction R package, with independent tools, reproducible workflows, visualization, and novel statistics.

## 1 Introduction

Combining genomic datasets from multiple studies increases statistical power in studies where logistical considerations limit sample size or require the sequential generation of data^4,5^. However, significant non-biological technical variation, or *batch effects*, frequently plagues high-throughput biological datasets^6^. This technical bias often arises from differences in reagents, experimental or lab protocols, or instrument platforms or settings^8^. Bias due to batch effects often confounds true biological relationships in the data, reducing the power benefits of combining multiple batches, and may even lead to spurious results if left uncorrected^9^.

Many R Shiny^10^ apps for batch effect correction have been created to streamline the data exploration, analysis, and visualization steps that accompany batch effect correction analyses, including our original beta version of BatchQC^1^ (Shiny app only), BatchFLEX^2^, and BatchServer^3^. Other apps, like DEBrowser^7^, also contain methods that address batch correction, while the primary use focuses on differential gene and region expression analyses. These apps enhance data exploration and visualization when determining the appropriateness of applying a batch correction method to a dataset by providing different batch effect correction methods (e.g. variations of ComBat^8^, Harman^11^, limma^12^), multiple visualization methods (e.g. heatmaps, PCA, UMAP), and other features (e.g. differential expression analysis, variation analysis) to aid in data exploration and downstream analyses (See Table 1; note that blue bold BatchQC features are introduced and detailed later in this manuscript).

**Table 1.** Feature comparison of various batch correction R Shiny apps. Methods and features highlighted in blue are introduced in BatchQC in this manuscript.

|  | BatchQC<br>Shiny<sup>1</sup> | Batch<br>FLEX<sup>2</sup> | Batch<br>Server<sup>3</sup> | DE<br>Browser<sup>7</sup> |
| --- | --- | --- | --- | --- |
| **Batch Corr. Methods** |  |  |  |  |
| autoComBat |  |  | x |  |
| ComBat | x | x |  | x |
| ComBat-Seq | **x** | x |  |  |
| Harman | **x** | x |  | x |
| limma | **x** | x |  |  |
| RUVg |  | x |  |  |
| sva | x | x | x |  |
| svaseq | **x** |  |  |  |
| **Visualizations** |  |  |  |  |
| All2all scatter plots |  |  |  | x |
| Circular Dendrogram | x |  |  |  |
| Dendrogram | **x** |  |  |  |
| Heatmaps | x | x |  | x |
| IQR and Density Plots |  |  |  | x |
| PCA | x | x |  | x |
| PCVA |  | x | x |  |
| Scree Plot |  | x |  |  |
| UMAP | **x** | x | x |  |
| Volcano Plot | **x** |  |  | x |
| **Evaluations** |  |  |  |  |
| Differential Expression | x |  |  | x |
| Goodness-of-Fit | **x** |  |  |  |
| kBET | **x** |  |  |  |
| lambda Statistic | **x** |  |  |  |
| Variation Analysis | x |  | x |  |
| **Additional Features** |  |  |  |  |
| CRAN/Bioconductor | x |  |  | x |
| Export Corrected Data | **x** | x | x |  |
| Normalization | **x** | x |  | x |
| Shiny Interface | x | x | x | x |
| Standalone R Functions | **x** | x |  |  |

A key feature that these shiny apps lack is a CRAN or Bioconductor R package with standalone R functions that can be utilized independently of the shiny exploration features. Most notably, R packages allow for greater reproducibility in analyses by enabling users to show the order of function calls, share analysis code and specific function parameters, and provide pre- or post-processing to datasets. Both DEBrowser^7^ and beta BatchQC^1^ are available on Bioconductor, but have previously only functioned as Shiny apps. BatchServer^3^ is a web-based shiny app with accompanying code hosted on GitHub. However, it does not have a companion R package to run functions independently of the web-hosted Shiny app. BatchFLEX^2^ is a web-based shiny app with a companion R package on GitHub, but it lacks documentation on how the shiny app’s analyses link to the companion package, which makes reproducibility more difficult. Additionally, these packages have focused on incorporating batch correction methods and visualization analyses without also incorporating quantitative methods for batch correction exploration and comparison.

In this paper, we present an updated BatchQC, a complete Bioconductor package featuring an object-oriented approach and an enhanced R Shiny app. As an R package, BatchQC analyses are completely reproducible and can be incorporated into complete analysis pipelines independent of the Shiny utility. In addition, all corrected datasets can be imported in and exported out of both the package and the Shiny app as a SummarizedExperiment object and easily incorporated into further downstream analyses. BatchQC contains both commonly used batch correction analyses and novel methods that enhance data exploration and analysis, including visual and quantitative approaches. Overall, BatchQC is a comprehensive R package that enables robust, reproducible batch correction analysis for a wide range of cases.

## 2 Methods

BatchQC utilizes a range of common batch effect evaluation tools (variation analysis, kBET^13^, dimension reduction, clustering, differential expression), as well as novel methods which are described in detail below. We developed a statistic, denoted λ, that measures batch signal-to-noise ratio and enables users to analyze the effectiveness of batch correction methods. Additionally, we created multiple model fit tools to suggest the best conforming model for downstream analysis tools. The utility of these novel methods was evaluated on simulation data as well as two real datasets. Detailed methods for each analysis tool, along with information about the datasets, are provided below.

### 2.1 Data Examples

Data for testing and validating our tools have been obtained from simulated data, a Tuberculosis (TB) gene expression study, and an Alzheimer’s imaging study (summarized in Table 2).

**Table 2:** Summary of the different datasets used throughout the paper to validate new methods and demonstrate BatchQC.

| Dataset Name | Dataset Source | Known Data Distribution | Use Case |
| --- | --- | --- | --- |
| Simulated Dataset 1 | simulated based on normal distribution | normal | $\lambda$ statistic validation for balanced and unbalanced designs |
| Simulated Dataset 2 | simulated from MOMS-PI data in HMP2Data package | negative binomial | AIC negative binomial validation |
| Simulated Dataset 3 | simulated from MOMS-PI data in HMP2Data package | log-normal | AIC log-normal validation |
| Simulated Dataset 4 | simulated from MOMS-PI data in HMP2Data package | voom-based | AIC voom validation |
| Simulated Dataset 5 | simulated from TCGA and GTEx | negative binomial | Goodness-of-Fit negative binomial validation |
| TB Dataset | RePORT Indian (Tuberculosis) TB/latent TB; whole blood gene expression | unknown | Demonstrate BatchQC tools |
| ADNI Dataset | Alzheimer's Disease Neuroimaging Initiative (ADNI); cortical thickness values | unknown | Demonstrate BatchQC tools |

#### Code Availability

All data, cleaning, and preprocessing steps are available on GitHub (github.com/jess-mcc22/BatchQCv2_Manuscript) except for the data needed for Simulation Dataset 5, which is available on Zenodo (https://doi.org/10.5281/zenodo.17429046).

#### Simulation Data

##### Simulation Dataset 1

We created a series of simulated datasets based on normal distributions to examine the impacts of batch effects, group effects, residual variations, and design balance on the false positive rate (FPR) and true positive rate (TPR) in differential expression analysis. Particularly, we designed three scenarios for group-batch design, namely the balanced design (5:5 ratio between the treatment and control groups across two batches), the unbalanced design (3:7 and 7:3 ratios between the treatment and control groups across two batches), and the extremely unbalanced design (1:9 and 9:1 ratios between the treatment and control groups across two batches). In addition, we also explored the balanced design (having the same number of participants from each batch) with unequal group sizes (i.e., 1:9, 2:8, 3:7, and 4:6) to ensure the robustness of our results. A range of variances for batch, group, and residual variations was utilized to yield diverse signal-to-noise ratios.

##### Simulation Dataset 2

We created a simulated dataset based on the MOMS-PI dataset from the Bioconductor HMP2Data package. An artificial batch effect was created via random assignment and the batch and condition variables were assigned to selectively create taxa with an associated effect based on the batch/condition variables. This enables us to create a dataset from a known negative binomial distribution, both with and without a batch effect.

##### Simulation Dataset 3

We created a simulated dataset based on the same metadata (group-batch design) of the MOMS-PI dataset from the Bioconductor HMP2Data package. The gene expressions were simulated using a log-normal distribution, with optional artificial batch effects and group/condition effects. This allows us to generate a dataset from a known log-normal distribution, both with and without batch effects.

##### Simulation Dataset 4

We created a simulated dataset based on the same metadata (group-batch design) of the MOMS-PI dataset from the Bioconductor HMP2Data package. The gene expressions were simulated using identical random-value-generator described in the Law et all.^14^ with optional artificial batch effects and group/condition effects. This allows us to generate a dataset from a known voom-based distribution, both with and without batch effects.

##### Simulated Dataset 5

A simulated dataset was created based on data available in The Cancer Genome Atlas Program (TCGA) and the Genotype-Tissue Expression (GTEx) Portal. For TCGA data, we selected RNA-seq datasets from six cancer types that have paired normal tissues and sample sizes greater than 50 for both cancer and normal tissues. Then, we downloaded the gene read count matrices of these selected datasets from GDC Xena Hub. For GTEx data, we selected six pairs of tissues with sample sizes ranging from 126 to 706.

We then downloaded the gene read count matrices for these tissue samples from the GTEx Portal. The synthetic datasets are generated as follows: for each gene, we generate counts from negative binomial distributions, where the mean for each sample and the dispersion parameter for this gene are both estimated from the negative binomial models of edgeR^15^. We generated one synthetic dataset for each GTEx and TCGA dataset, totaling 12 synthetic datasets.

#### Real-World Datasets

##### Tuberculosis Study

The TB study contains whole blood gene expression profiling from well and malnourished Indian individuals with TB and severely malnourished household contacts with latent TB, which were originally obtained through the Regional Prospective Observational Research in TB (RePORT) study based at Jawaharlal Institute of Postgraduate Medical Education and Research (JIPMER). The data was originally obtained in two different studies and merged to create a larger sample size^16^.

##### Alzheimer’s Study

The Alzheimer’s study is a dataset obtained from the Alzheimer’s Disease Neuroimaging Initiative (ADNI) database (adni.loni.usc.edu). The ADNI was launched in 2003 as a public-private partnership, led by Principal Investigator Michael W. Weiner, MD. The primary goal of ADNI has been to test whether serial magnetic resonance imaging (MRI), positron emission tomography (PET), other biological markers, and clinical and neuropsychological assessment can be combined to measure the progression of mild cognitive impairment (MCI) and early Alzheimer’s disease (AD). For up-to-date information, see www.adni-info.org.

The ADNI dataset used in this manuscript contains cortical thickness values of various locations in the brain in a longitudinal study obtained from the ADNI database in May 2014, as described in Tustison et al.^17^ Scans were taken on a Siemens Healthineers scanner or a Phillips Medical System/GE scanner. Our analysis focuses on cortical thickness scans of the temporal region of the brain taken at the first visit. Previous work indicated a batch effect in this data set when using DeepCombat^18^, which is specifically designed for imaging data. We show in this manuscript that this batch effect can be identified with the tools in BatchQC.

### 2.2 Object-Oriented Design

The object-oriented interface of BatchQC enhances users’ experience by allowing greater flexibility in data upload and download, analysis, and reproducibility. Users may load their data as a SummarizedExperiment (SE) object^19^ (which is the standard and recommended data structure for genomics data from Bioconductor^20^) or as two matrices, one containing the feature data and the other containing the metadata. Once data is loaded, users can then choose the individual analyses, normalization, and correction methods to apply to their data. After exploration and analysis, users may export an SE object that contains all corrected matrices, as well as the original data uploaded (Table 1). As a standard Bioconductor data structure, the exportable SE object can be easily utilized with other downstream Bioconductor packages, such as DESeq2^21^, edgeR^15^, or MultiAssayExperiment^22^, among others. Additionally, many parameters within the BatchQC analyses can be set and adjusted by the user, allowing for more fine-tuned data exploration.

BatchQC is a complete R package that enables command-line usability and integration into analysis pipelines (Table 1). To enhance data exploration and provide a beginner-friendly interface, the BatchQC package also includes an R Shiny application that allows point-and-click data upload, exploration, and download (Table 1).

Vignettes are available detailing the example datasets (“BatchQC Examples”) available in BatchQC and an example workflow (“BatchQC Intro”) for the Shiny app and the accompanying command-line code. The “BatchQC Examples” vignette is available at https://bioconductor.org/packages/3.23/bioc/vignettes/BatchQC/inst/doc/BatchQC_examples.html and the “BatchQC Intro” vignette is available at https://bioconductor.posit.co/packages/3.23/bioc/vignettes/BatchQC/inst/doc/BatchQC_Intro.html.

### 2.3 Common Batch Assessment Methods Normalization

We have implemented common normalization methods utilized in RNA-Seq data analysis, including counts per million, log, DESeq2^21^, edgeR^15^, and voom^14^ normalizations. Users can specify the SummarizedExperiment object, the assay they would like to normalize, and a name for the created normalized assay; the normalized assay is then added to the SE object. This feature can be accessed on the “Batch Correction/Normalization” tab in the shiny app or via the “normalize_SE()” function in the package.

#### Correction Methods

Users can apply one of many batch correction methods to an assay contained within their selected SummarizedExperiment object. The corrected assay is provided as output as an assay in the SummarizedExperiment, with a name specified by the user. Users may select from the following correction methods, all of which are implemented according to the package specifications pertaining to the package: ComBat^8^, Combat-Seq^23^, Harman ^11^, limma^12^, sva^24^, and svaseq^25^. This feature can be accessed on the “Batch Correction/Normalization” tab in the shiny app or via the “batch_correct()” function in the package.

#### Experimental Design

Users can provide their batch and biological variables (in both the shiny app on the “Experimental Design” tab with the subtab “Batch Design” and via the “batch_design()” function) and receive output displaying the number of samples in each batch and each biological variable. Confounding statistics (Pearson correlation coefficient and Cramer’s V estimation) are also provided based on the selected batch variable (in the shiny app under the “Experimental Design” tab with the “Confounding Statistics” subtab or via the “confound_metrics()” function). The Pearson correlation coefficient indicates the strength of the linear relationship between the batch and condition variables. Note that this is the correlation between two categorical variables, so rather than measuring the correlation between the two variables (as with two continuous variables), it directly measures the dependence/independence of the batch and condition in the experimental design—i.e., the potential for confounding, but not confounding itself. Values for this metric can range from -1 to 1, with a value close to 0 indicating batch/condition independence or a “balanced design”, which should be associated with a lower risk of observing a batch effect. A value closer to -1 or 1 indicates greater imbalance in the design and a higher likelihood of a batch effect. The Cramer’s V provides a similar metric (ranging from 0 to 1) for experimental design balance, with values closer to 0 indicating a more balanced design and a lower likelihood of a batch effect, and values closer to 1 are associated with an imbalanced design and a greater likelihood of a batch effect.

#### Variation Analysis

BatchQC provides information on the individual (marginal) variation explained by each variable independently, as well as the Type II partial variation (based on Type II sums of squares), which represents the variance explained by a factor after accounting for other main effects in the model. It also provides ratios displaying the individual variation/batch ratio and residual variation/batch ratio. For each type of variation, a ggplot2^26^ boxplot is displayed, along with a table listing the variation specific to each gene. This table is searchable and can be displayed in ascending or descending order. The larger the amount of explained variation attributed to the batch variable, the more likely there is a batch effect in the dataset. For “Individual Variation Variable/Batch Ratio” and “Residual Variation Variable/Batch Ratio,” a ratio greater than 1 indicates that the batch has a stronger effect than the variable of interest. When a batch effect is being properly corrected, the variation explained by the batch variable should decrease in the batch-corrected assay, indicating that the variation in the data is now more likely to be associated with the condition of interest rather than the batch. The ratio statistics should also be less than 1. This functionality can be found in the shiny app under the “lambda/Variation Analysis” tab or by utilizing the “batchqc_explained_variation()” function and passing the output to “EV_plotter()” and/or “summary_stats_EV_table()” as described in the “BatchQC Intro” vignette.

#### kBET

The kBET algorithm used in BatchQC is from the Theis Lab^13^. kBET calculates an overall rejection rate by averaging a set of χ^2^-based binary tests to determine whether neighborhoods are well mixed; a lower rejection rate indicates well-mixed replicates, which in turn indicate a lower likelihood of a batch effect being present. BatchQC produces a boxplot of observed and expected rejection rates, as well as a table with the values and associated p-values comparing the observed and expected rates for the mean, quartile 1, the median, and quartile 3. To our knowledge, BatchQC is the first package or shiny app to implement kBET as part of a batch correction pipeline. The kBET functionality is available in the shiny app under the “kBET” tab or by utilizing the “run_kBET()” function and passing the output to “plot_kBET()”.

#### Clustering

##### Heatmaps

BatchQC includes a sample correlation heatmap and a gene-level heatmap comparing samples and genes; both are created using pheatmap^27^. The sample correlation heatmap shows how similar each sample is to each other sample in the data and the samples are clustered together based on similarity. The gene-level heatmap displays patterns of gene expression across the samples based on the selected variables of interest. These heatmaps are available on the “Heatmaps” tab, under their respective subtabs, in the shiny app, and can be created using the “heatmap_plotter()” function in the package.

##### Dendrograms

Dendrograms in BatchQC are created using ggdendro^28^, an extension of ggplot2^26^. Samples are clustered based on similarity of gene expression and other metadata using a Euclidean distance metric. The circular dendrogram displays the same information as the dendrogram, but in a circular format to reduce the distance between the top and bottom branches of the dendrogram. They can be accessed on the “Dendrograms” tab, under their respective subtabs” in the shiny app or by utilizing the “dendrogram_plotter()” function.

#### Dimension Reduction

Principal Component Analysis (PCA) is completed with the stats^29^ package and plotted using ggplot2^26^. Uniform Manifold Approximation and Projections (UMAP) are created using the umap^30^ R package. Both allow the user to compare how tightly clustered the batch variable is in comparison to the biological variable of interest. When the batch variable clusters more distinctly than the biological variable, there is likely a batch effect present. Within the shiny app, the PCA plots can be created under the “PCA” tab and the UMAP plots can be created under the “umap” tab. To use from the command line, “PCA_plotter()” will create a PCA plot and “umap()” will create a umap plot.

#### Differential Expression

BatchQC implements differential expression analyses from the following R packages: DESeq2^21^, limma^12^, edgeR^15^, and base R stats^29^ (for ANOVA and Kruskal-Wallis methods). DESeq2 should be used for count data (non-negative, integer values) with a negative binomial distribution, edgeR can be used for edgeR normalized data, ANOVA can be used for normally distributed data, Kruskal-Wallis test can be used as a nonparametric equivalent of one-way ANOVA, and limma can be used for all other data. The edgeR implementation in BatchQC uses glm quasi-likelihood F-tests to test for differential expression. Volcano plots based on the differential expression analysis output are generated using ggplot2^26^. This functionality can be found under the “Differential Expression” shiny app tab with subtabs for the “Results,” “p-value analysis,” and “Volcano Plot.” To complete on the command line, run the “DE_analyze()” function to view the results (as seen on the “Results” tab), pass the results to “pval_plotter()” and “pval_summary()” to recreate the “p-value analysis” subtab, and “volcano_plot()” to recreate the “Volcano Plot” tab. More details and examples are provided in the BatchQC Intro Vignette.

### 2.4 Novel Batch Effect Evaluation Metrics Signal-to-noise evaluation (Lambda Statistic)

We have developed a metric, henceforth referred to as our 𝜆 statistic, based on the biological signal to technical (batch) noise ratio that can be used to quantify the impact of a batch effect. Specifically, it is defined as follows:

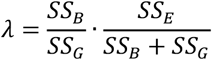

where 𝑆𝑆_𝐵_, 𝑆𝑆_𝐺_ and 𝑆𝑆_𝐸_ denote the sum of squares due to the batch effect, the group effect, and the error (residuals). To gain intuition about our 𝜆 statistic, one can perceive the above definition as the ratio between the relative size of the batch effect (which equals to 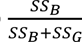) and the relative size of the group effect (which equals to 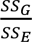). It is worth noting here that the relative size of the batch effect quantifies the importance of the batch effect, relative to the biological signal, while the relative size of the group effect characterizes the importance of the biological signal, compared to noise. According to the definition, one would expect our 𝜆 statistic to increase if the relative size of the batch effect increases or the relative size of the group effect decreases, indicating that the necessity of batch effect adjustment increases. We have determined that a λ statistic greater than -7 is required to preserve a statistically valid false-positive rate in data sets with a balanced design. In experiments with an unbalanced design, a batch correction method should always be considered to preserve a statistically valid false-positive rate. This threshold for the 𝜆 statistic, including recommendations for both balanced and unbalanced designs, and performance is evaluated later in the results section using simulated dataset 1. This metric is available in the shiny app under the “lambda/Variation Analysis” tab and can be obtained with the “lambda_stat()” function.

#### Distribution Goodness-of-Fit

We created two separate tools to assess the model fit of a given dataset. The first approach suggests the best overall model fit for a dataset between the negative binomial, log-normal, and voom variance stabilizing transformation (Gaussian-like) distributions. The second method directly tests the goodness-of-fit (GoF) for a negative binomial assumption, such as is needed for ComBat-Seq batch effect correction or differential expression using DESeq2^21^ or edgeR^15^. We propose separate approaches for negative binomial GoF on both small and large datasets. We incorporate the GoF comparison and model selection index to assist users in selecting the most suitable model for RNA-seq data. Both methods can be accessed on the shiny app under the “Batch Correction/Normalization” tab and their respective subtabs, “Negative Binomial Check” or “AIC Computation.” Run “compute_aic()” for the AIC output and “goodness_of_fit_nb()” for the GoF for negative binomial.

#### AIC Goodness-of-Fit

The Akaike Information Criterion (AIC) was utilized to compare the model fits of negative binomial (ComBat-seq), log-normal (ComBat) and voom variance stabilizing transformation (Gaussian-like) distributions^14^ (limma), to the data in order to determine the appropriateness of ComBat-seq, ComBat, and voom/limma, respectively, in batch correction, as these methods are all parametric in nature with rigid distributional assumptions. We provide an AIC score to estimate the best distributional fit for the data that is a ratio of the total AIC to the minimum AIC. These are calculated as follows:

Suppose we have 𝐾 different models in comparison (we are using 3 models in comparison, negative binomial, log-normal and voom), then the total AIC for the 𝑖𝑡ℎ model (𝑖=1,⋯,𝐾) is defined as:

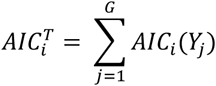

where 𝐴𝐼𝐶i(𝑌j) denotes the AIC value calculated based on 𝑌j, which is a vector of the 𝑗^th^ gene’s expressions across all samples, under the 𝑖^th^ model, assuming the total number of genes is 𝐺. The minimum AIC is defined as:

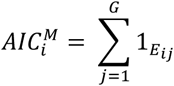

where the event 𝐸ij is defined as the event that 𝐴𝐼𝐶i(𝑌j) is the minimum value across all 𝐾 different models for the 𝑗^th^ gene, i.e.:

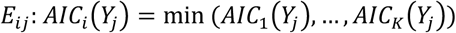

The AIC score (𝛽) is then defined as:

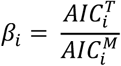

By accounting for the minimum AIC per gene (𝐴𝐼𝐶^𝑀^), we reduce the excessive influence of a few outlier genes on the AIC score. We can compare AIC scores (𝛽i) across different models; the optimal model should possess the minimum 𝛽i value. We also calculate and report the mean AIC score for each method. Multiple simulation datasets (datasets 2, 3, and 4) were used to demonstrate that this method accurately identified the correct distribution of the provided data and are presented in the results section.

#### Negative Binomial Goodness-of-Fit

We assess the negative binomial (NB) distribution assumption using a prior study that evaluated the appropriateness of distributional assumptions^31^, leveraging the edgeR package^15^. The prior work derived a model-free method for large datasets (more than 20 samples) and, in BatchQC, we provide the first software implementation. Additionally, we have derived a model-based method for analyzing 20 or fewer samples based on this work.

The model-based method employs a GoF comparison to evaluate whether the negative binomial model, commonly used in existing methods like DESeq2^21^ or edgeR^15^, is an appropriate fit for the data. The comparison involves the following steps (as adapted from previous work^31^):

1. For each gene under each condition, we use the unnormalized counts and the fitted NB model (from DESeq2^21^ or edgeR^15^). The model’s mean parameter is specific to each count, while the dispersion parameter is shared across counts under the same condition. We convert each count into a cumulative distribution function (CDF) value based on the fitted NB model. If the NB model fits well, these CDF values should be independent and uniformly distributed between 0 and 1.
2. To assess the uniformity of the CDF values, we perform a Kolmogorov–Smirnov (KS) test to evaluate how well a uniform distribution between 0 and 1 fits the CDF values. The resulting p-value reflects the GoF, with smaller p-values indicating a poorer fit.
3. We calculate the proportion of small p-values (e.g., p-values below 0.01). A dataset is considered to meet the NB model assumption if fewer than 1% of genes have p-values below 1%. This threshold is confirmed by analyses of datasets subsampled from the GTEx, TCGA, and synthetic datasets generated under the NB model (simulated dataset 5) as described in the results.

## 3 Results

### 3.1 BatchQC Overview

BatchQC is an R package with an R Shiny interface, built on an object-oriented framework (Table 1). It provides an enhanced batch correction analysis experience through a comprehensive R package, an R Shiny application, and the capability to export all data for downstream analysis. It is reproducible and can be seamlessly integrated into analysis pipelines.

Additionally, BatchQC includes multiple batch correction methods (including ComBat^8^, Combat-Seq^23^, Harman^11^, limma^12^, sva^24^, and svaseq^25^) and common batch effect analyses (such as variation confounding statistics, kBET, clustering, dimension reduction, and differential expression). Each of these tools enables users to explore the need to batch correct their raw data and/or evaluate the effectiveness of their chosen batch correction method on corrected data. Details regarding the implementation of each are described more fully in the methods. We have also included novel methods to aid batch correction analysis, which are detailed in the methods and evaluated below.

### 3.2 Novel Methods on Simulated Data

Here we evaluate performance of the new batch effect methods (see Methods for details), including our λ statistic based on variation analysis, our distribution goodness-of-fit (GoF) models, an AIC comparison and an edgeR^15^-based negative binomial GoF. Each method was tested on simulated data (see Table 2) and the results are provided below.

#### Variation Analysis and λ Statistic

BatchQC evaluates both individual and residual (Type II) variation. Additionally, individual variation/batch and residual variation/batch ratios are also provided, where ratios greater than 1 indicate that the variation present in the batch variable is greater than in the experimental variable. The results for individual, residual, and both ratios are presented as boxplots.

The intuition for our λ statistic is based on the signal-to-noise ratio for the variable of interest. When the signal-to-noise ratio is small, a batch correction is unlikely to be needed, whereas when the signal-to-noise ratio is large, batch correction is likely beneficial.

We validated our λ statistic utilizing dataset 1 with a combination of balanced and unbalanced designs as described in the methods (Table 2). We assessed how the true positive rate (TPR) and the false positive rate (FPR) for the benchmark (no batch effect), unadjusted (with batch effect), and ComBat-corrected datasets changed based on the variation represented by the λ statistic. In datasets with perfectly balanced designs (5:5), the unadjusted TPR fell below the benchmark and ComBat corrected TPR when the λ value was greater than -7 (Figure 1A). Therefore, we recommend that perfectly balanced designs with a λ statistic greater than -7 should be batch-corrected to conserve TPR. However, in the unbalanced (Figure 1B) and extremely unbalanced (Figure 1C) simulated datasets, the unadjusted FPR is consistently above 0.05 (about 0.01 in Unbalanced designs regardless of λ statistic and above 0.25 in Extremely Unbalanced Designs across the λ statistic value). Therefore, batch correction methods should always be applied to preserve a statistically valid false positive rate in datasets with unbalanced designs. In balanced designs with unequal ratios in the batches, as the ratio becomes more unbalanced, batch correction aids in maintaining a statistically valid FPR (Supplemental Figure 1). Applying a batch correction in these balanced, but unequal batch distribution ratio scenarios also maintains a TPR that is more similar to the unadjusted TPR. Therefore, as the distribution of samples between batches becomes more unbalanced, or as a balanced design approaches a λ greater than -7, it is likely best to apply a batch correction method to preserve TPR and maintain a statistically valid FPR.

**Figure 1:**
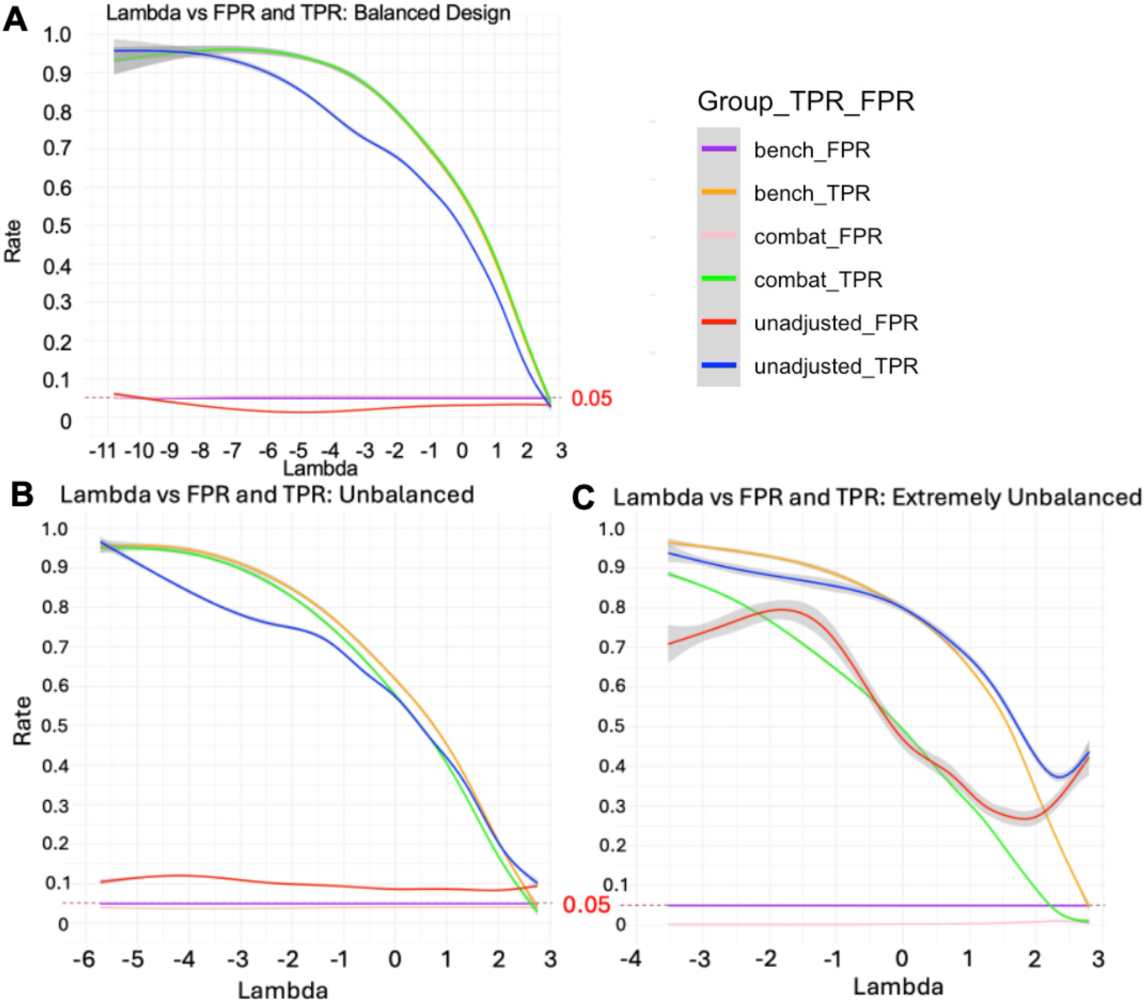
False positive rates (FPR) and true positive rates (TPR) based on experimental design. FPR (red) and TPR (blue) before batch correction, after ComBat batch correction (FPR is pink, TPR is green), and for benchmark data (FPR is purple and TPR is orange) for A) Balanced experimental design, B) Unbalanced experimental design, and C) Extremely unbalanced experimental design. To preserve TPR and reduce FPR, correction methods should be applied to balanced designs only when the variance explained by the batch variable exceeds a lambda value of -7 (A). However, correction methods should always be applied to unbalanced designs (B and C) to preserve the false positive rate.

#### Akaike Information Criterion Goodness-of-Fit

The Akaike Information Criterion (AIC) GoF compares AIC metrics for negative binomial, log-normal, and voom/limma Gaussian distributions on the sum of the AIC scores for the dataset as a whole (total AIC), on the count of the number of genes with the lowest AIC for each individual gene within a dataset (minimum AIC), the median AIC for each method (median AIC), and as the ratio between the total AIC and the minimum AIC (AIC score). The best-fitting distribution should have the minimum AIC score. We applied the AIC-based score to multiple simulation studies (Table 2), including datasets generated directly from negative binomial distribution, log-normal distribution, as well as voom-based distribution, to confirm that the AIC score can identify the correct distribution (Figure 2).

**Figure 2:**
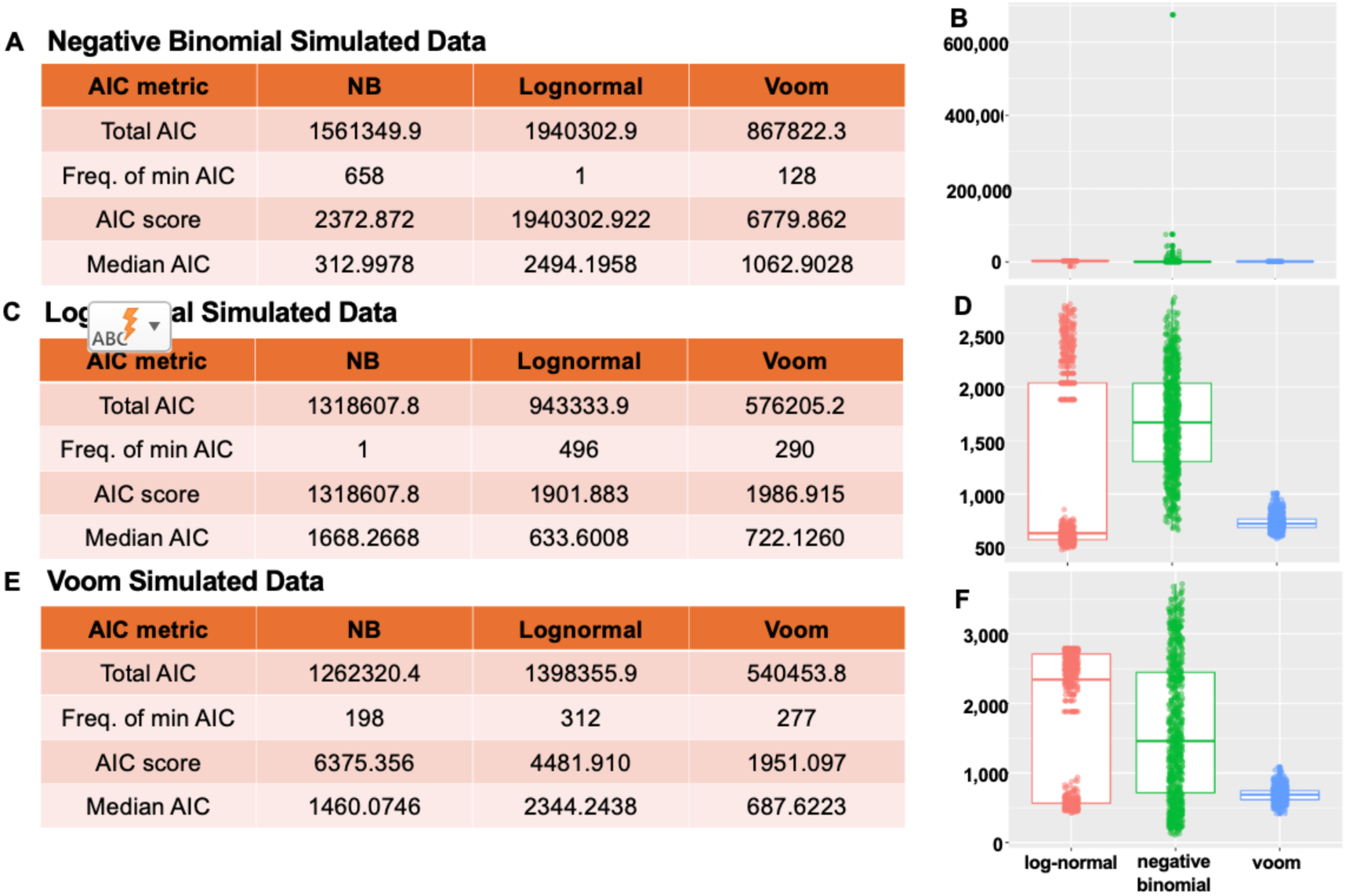
Akaike Information Criterion (AIC) model fit on simulated datasets. The AIC score correctly predicts the model fit of the model created from the simulated data. A) Negative binomial (NB) simulated data has the lowest AIC score for the NB model. B) Distribution of individual AIC scores for each gene in each model for the NB simulated data. C) Log-normal data has the lowest AIC score for the log-normal model. D) Distribution of individual AIC scores for each gene in each model for the log-normal simulated data. E) Voom simulated data has the lowest AIC score for the voom model. F) Distribution of individual AIC scores for each gene in each model for the voom simulated data.

For simulated dataset 2 with a negative binomial distribution, the AIC scores were 2372.872 for negative binomial, 1940302.922 for log-normal, and 6779.862 for voom (Figure 2A). Since 2372.872 is the lowest score, the data seems to best fit a negative binomial distribution, as expected. For simulated dataset 3 with a log-normal distribution, the AIC scores were 1318607.8 for negative binomial, 1901.883 for log-normal. and 1986.915 for voom (Figure 2B). As expected, the log-normal score (1901.883) is the lowest, which would predict the log-normal distribution to be the best fit for the data. Finally, for simulated dataset 4 with a normal distribution, we would expect the lowest AIC score for voom. The results support a normal distribution with AIC scores of 6375.356 for negative binomial, 4481.910 for log-normal, and 1915.097 for voom (Figure 2C).

Furthermore, when ComBat, ComBat-Seq, and limma are applied as correction methods on each of the simulated datasets (2, 3, and 4), the results for the method that requires the specific distribution simulated in the datasets mimic those found in the raw data without a batch effect (See Supplemental Figures 2-4). This indicates that the AIC score not only correctly predicts the dataset’s distribution but, when used with appropriate downstream tools, yields accurate results.

#### Negative Binomial Goodness-of-Fit

As detailed in the methods, we created a model-based test based on edgeR for datasets with less than or equal to 20 samples and utilized the previously described model-free test^31^ that can be applied to datasets with larger sample sizes. We evaluated the performance of the model-based method using synthetic dataset 5 (Table 2).

Specifically, we subsampled both the original GTEx/TCGA datasets and the corresponding synthetic datasets to create small-sample datasets of 10 and 20 samples and then applied the model-based method to obtain a p-value for each gene in each dataset. We computed the proportion of p-values below 1% across all genes. The results show that for datasets subsampled from the original data, which are likely affected by model misspecification, the proportion of small p-values consistently exceeded 1%. Specifically, in the original data with 10 samples, the minimum value was 0.0109 and for the original data with 20 samples, the minimum value was 0.0257 (Figure 3A). In contrast, for datasets subsampled from the synthetic data, the proportion of small p-values was mostly below 1% (maximum = 0.003 in 10 samples and maximum = 0.007 with 20 samples; Figure 3B). Based on these observations, we selected 1% as the threshold for the model-based method.

**Figure 3:**
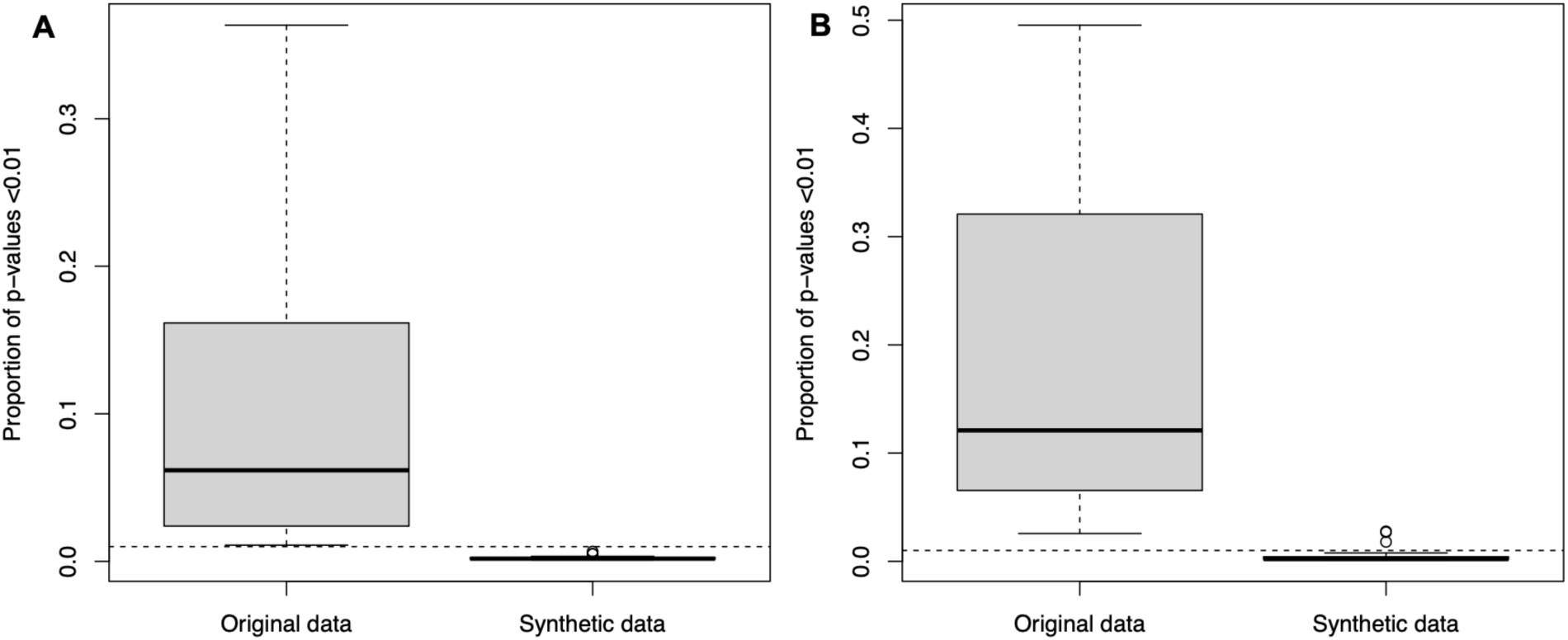
Boxplot of p-values for negative binomial Goodness-of-Fit simulated results. A) Proportion of p-values less than 0.01 for sample size of 10 (5 per condition) and B) when sample size is 20 (10 per condition). For datasets subsampled from the original data (likely affected by model misspecification), the proportion of small p-values consistently exceeded 1% (the dotted line; minimum = 0.0109 for the original data in A and minimum = 0.0257 for the original data in B), whereas for datasets subsampled from the synthetic data, the proportion of small p-values was mostly below 1% (maximum = 0.003 in synthetic data in A and maximum = 0.007 in synthetic data in B) Based on these observations, we selected 1% as the threshold for the model-based negative binomial goodness-of-fit.

When the proportion of significant p-values is less than the threshold (0.01 for model-based, and the number of genes/1000 for the model-free approach), the data likely conforms to a negative binomial distribution.

### 3.3 Real Data Applications TB Data

#### Determine Need for Batch Correction

The TB dataset is provided as an example dataset in the BatchQC package and can be loaded from within the package. After uploading the data, we first assess the need for batch correction through variation (including λ statistic), PCA, UMAP, and kBET analyses.

In the variation analysis, the batch variable, “Experiment”, shows a greater amount (p < 0.001) of explained variation (mean = 15.8) than the biological variable, “TB Status” (mean = 5.5), in the residual variation analysis (Figure 4A). Since this dataset has an unbalanced design, our λ-statistic analysis indicates that we should apply a batch correction method to preserve the false positive rate, regardless of the λ statistic (which was 3.14). Furthermore, in both the PCA (Figure 4E) and UMAP (Figure 4I) plots, a clear separation is observed between the different batch variables. Finally, in the kBET analysis (Figure 4M), the observed rejection rate (mean = 0.866) is significantly (p = 0) greater than the expected rejection rate (mean = 0), another indicator that this dataset would likely benefit from a batch correction analysis.

**Figure 4:**
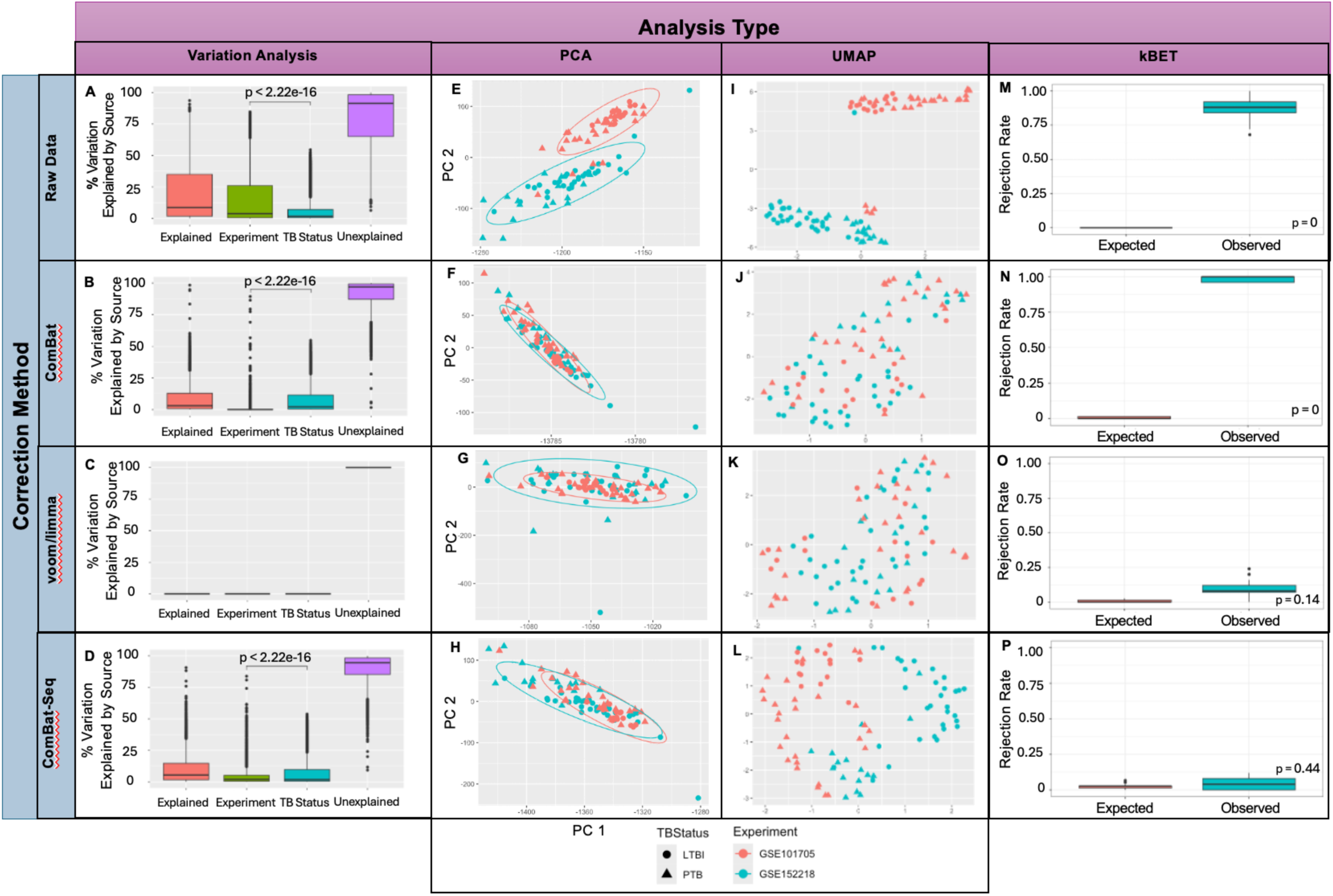
Raw and corrected Tuberculosis (TB) data batch analysis. Residual variation analysis: A) Raw, uncorrected data. The batch variable, “Experiment” (mean = 15.8), explains more variation (p < 0.001) than the biological variable, “TB Status” (mean 5.6), indicating that a batch effect is present. B) ComBat corrected data where the biological variable (mean = 7.3) accounts for most of the explained variation (p <0.001), rather than the batch variable (mean = 0.4). This indicates that ComBat is properly adjusting for the batch variable. C) voom/limma corrected data where there is no explained variation due to either the batch (mean = 0) or biological variable (mean = 0), indicating overcorrecting the data (no p-value to report). D) Combat-Seq reduces the amount of variation (p < 0.001) attributed to the batch variable (mean = 3.5) compared to the biological variable (mean = 6.9), making it a potential batch correction method. **Principal component analysis (PCA):** E) Raw, uncorrected data shows almost complete separation between the two batches (Experiment variable), rather than by TBStatus (Latent TB Infection (LTBI) and Positive TB (PTB)), indicating a need for batch correction. F) ComBat corrected data shows homogeneous mixing between the batches, an indicator that ComBat may be a good batch correction method to utilize. G) voom/limma has one batch (GSE101705) completely contained within the other batch, a sign that overcorrection may be occurring. H) ComBat-Seq has more homogenous mixing compared to the raw data and may be an appropriate correction method to select. **Unifold Manifold Approximation and Projection (UMAP) analysis:** I) Raw, uncorrected data shows two distinct groups for the batch (Experiment) variable, indicating a need for batch correction. J) ComBat shows homogenous mixing of the batch variable with stronger grouping by the biological (TB Status) variable. K) voom/limma shows more homogenous mixing of both the batch and the biological variable, a sign of over correction. L) ComBat-Seq still shows separation between the batch variable. **kBET analysis:** M) Raw data has a mean observed rejection rate that is significantly higher than the mean expected rejection rate (p = 0), indicating a strong batch effect in the TB dataset. N) ComBat corrected data still has a mean observed rejection rate that is significantly higher than the mean expected rejection rate (p=0), indicating that the batch effect is still present. O) voom/limma corrected data has a mean observed correction rate that is more similar to the mean expected rejection rate (p = 0.14), indicating that the batch effect has been corrected. P) ComBat-Seq also has a mean observed correction rate that is more similar to the mean expected rejection rate (p = 0.44), indicating that the batch effect has been corrected.

#### Batch Correction

After determining a need to apply a batch correction method, we can use our GoF AIC comparison and negative binomial GoF to estimate the distribution of the dataset to aid our decision on which correction and other downstream methods would be appropriate for our dataset. We can also apply any needed normalizations prior to batch correction.

The AIC score is lowest for log-normal (331.9807), followed by voom (977.6471), and the negative binomial distribution (18716166) (see Supplemental Figure 5). This suggests that we should consider using tools that assume a log-normal distribution, such as ComBat. We can further evaluate this by applying our negative binomial GoF procedure, which found that 36% of our features were significant, supporting the hypothesis that a negative binomial distribution is unlikely to be a good fit for the data. We will further evaluate the effectiveness of ComBat, voom/limma, and ComBat-Seq, all of which are common batch correction methods for RNA-Seq data, but anticipate ComBat being the best method for batch correction.

We can normalize our data with a counts per million log transformation before applying ComBat and apply a voom normalization prior to using limma. ComBat-Seq can be applied directly to the counts data.

#### Analyze Batch Correction Methods

We first compare the variation analysis of each correction method to the raw data. In the ComBat variation analysis (Figure 4B), most of the explained variation (p<0.001) is attributed to the biological variable, "TB Status” (mean = 7.3), and the batch variable, “Experiment” (mean = 0.4), no longer explains much of the variation. This indicates that ComBat is properly adjusting for the batch variable while preserving the biological variable. The voom/limma variation analysis shows signs of overfitting as the explained variation is no longer attributed to either the batch (mean = 0) or the biological (mean = 0) variable (Figure 4C). ComBat-Seq shows improvement over the raw data; however, the batch variable (mean = 3.5) still explains some of the remaining variation, although significantly less (p < 0.001) than the biological variable (mean = 6.9) (Figure 4D). When we calculate a λ statistic for each method and compare it to the raw λ of 3.14, we see that each method improves the overall batch effect (ComBat λ = -0.62, voom/limma λ = Inf, and ComBat-Seq λ = 2.55). However, λ = Inf is another sign of overcorrection for voom/limma and ComBat-seq only shows slight improvement.

Next, we compare the PCA and the UMAP for each method. ComBat shows homogenous mixing between the batch variable in both the PCA (Figure 4F) and the UMAP (Figure 4J) plots. Meanwhile, voom/limma again shows signs of overcorrection as one batch (GSE101705) is completely contained within the other batch (GSE152218) in the PCA plot (Figure 4G). We do observe homogeneous mixing of both the batch and biological variables in the voom/limma UMAP (Figure 4K). For ComBat-Seq, we observe a more homogeneous mixing of the batch variable in the PCA plot (Figure 4H), although some separation of the batch variables remains in the UMAP (Figure 4L). These analyses suggest that ComBat would be a suitable correction method.

Finally, we compare kBET analyses. ComBat shows an increased observed rejection rate (mean = 0.986) compared to the raw data (mean = 0.866) and is still significantly (p = 0) different from the expected rate (mean = 0.004; Figure 4N). Meanwhile, both voom/limma (mean = 0.098; Figure 4O) and ComBat-Seq (mean = 0.045; Figure 4P) show an improved observed rejection rate in comparison to the raw data (mean = 0.888) and both are not significantly different from the expected rejection rate (voom/limma p = 0.14 and mean = 0.006; ComBat-Seq p = 0.44 and mean = 0.021). Based on the kBET analysis, voom/limma or ComBat-Seq would be better correction methods to utilize than ComBat.

All but the kBET analyses indicated that ComBat would sufficiently correct the batch effect observed in the dataset. Therefore, we would perform a ComBat correction on the raw data and utilize the ComBat-corrected data in downstream analyses.

#### ADNI Data

##### Determine Need for Batch Correction

After loading the ADNI dataset, we first analyze the dataset for a batch effect through variation (including the λ statistic), PCA, UMAP, and kBET analyses.

In the variation analysis, we see that the batch variable, “Manufacturer” (mean = 7.4), accounts for a comparable amount of variation (p = 0.19) as the biological variable, “Diagnosis” (mean = 10.8), indicating the possibility of a batch effect. (Figure 5A). Furthermore, this dataset is unbalanced with a λ = 0.74. Therefore, to preserve the false positive rate, batch correction is recommended regardless of the value of λ. Additionally, in both the PCA (Figure 5E) and UMAP (Figure 5I) plots, we see that clustering occurs more strongly based on “Manufacturer” than on “Diagnosis,” again indicating the presence of a batch effect.

**Figure 5:**
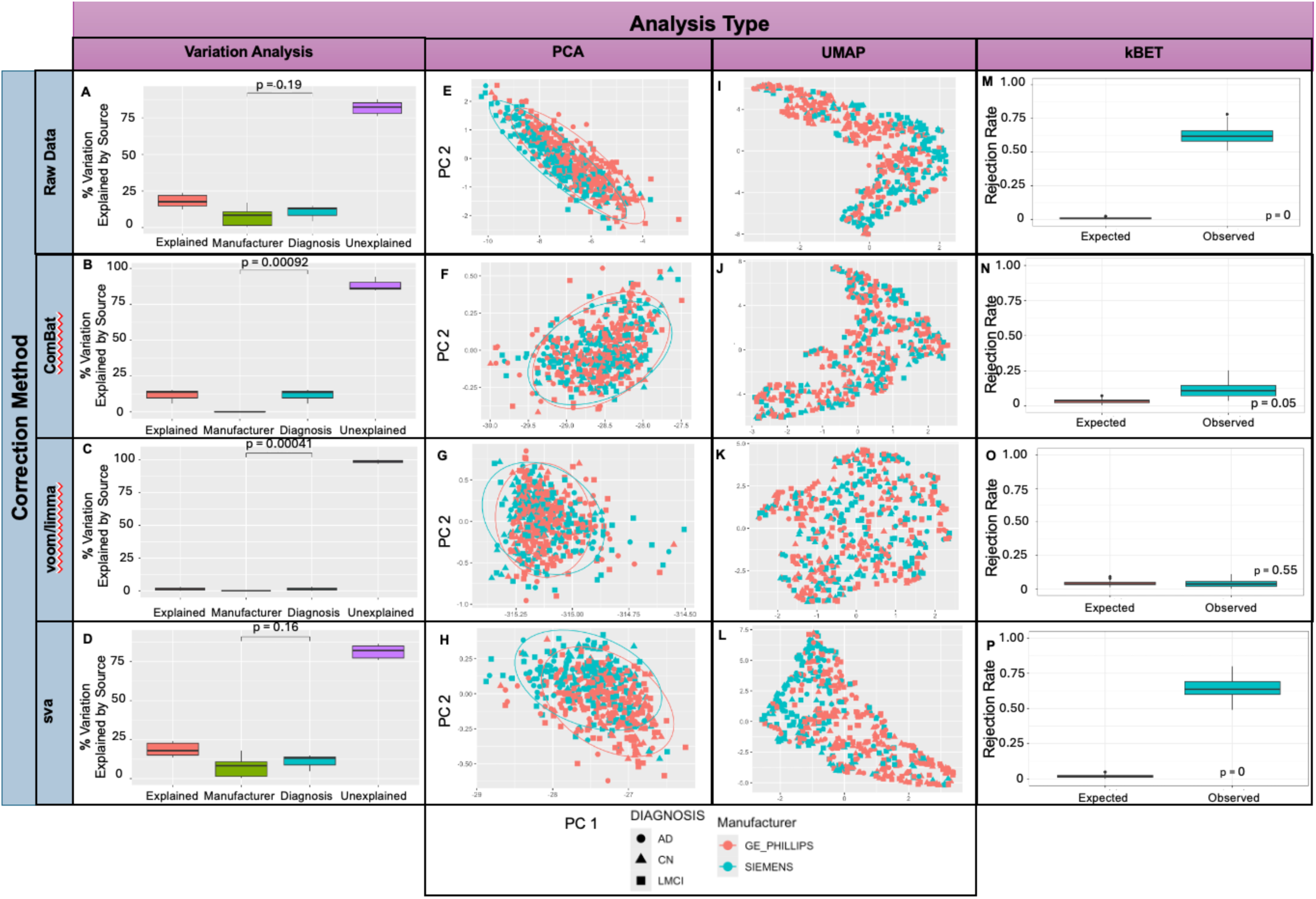
Raw and corrected ADNI data batch analysis. Residual variation analysis: A) Raw data shows similar variation (p = 0.19) explained by the batch (Manufacturer, mean = 7.4) and the biological (Diagnosis, mean = 10.8) variable, indicating a need for batch correction. B) ComBat corrected data shows little variation explained by the batch variable (mean = 0.04) compared to the biological variable (mean = 11.8), with a p < 0.001. Therefore, ComBat may be a good correction method to utilize. C) voom/limma correction shows little variation explained by the batch (mean = 0) or biological variable (mean = 1.29), an indicator of overcorrection (p <0.001). D) sva corrected data shows a similar amount of variation explained by the batch (mean = 7.4) and biological variable (mean = 11.1), indicating that sva does not properly correct for the batch effect (p = 0.16). **Principal component analysis (PCA):** E) The raw, uncorrected data shows some separation based on the batch variable (Manufacturer), rather than the biological variable (Disease; Alzheimer’s Disease (AD), Cognitive Normal (CN), and Late Mild Cognitive Impairment (LMCI)), indicating a need for batch correction. F) ComBat corrected data shows homogenous mixing between the batch variable, indicating a well-corrected dataset. G) voom/limma also shows homogenous mixing between the batch variables, indicating that the batch effect has been corrected. H) sva correction shows no change, indicating that a batch effect was not corrected. **Uniform Manifold Approximation and Projection (UMAP) analysis:** I) Raw Data shows some separation between the batch variable (Manufacturer), where the GE_PHILLIPS group is more to the upper left and the SIEMENS group is more to the lower left. This indicates a slight batch effect. J) ComBat shows homogeneous mixing between the batch variable, indicating that it would be a good correction method. K) voom/limma also shows homogenous mixing between the batch variable, indicating that also corrects for the batch effect. However, we also see more homogenous mixing of the biological variable (Diagnosis), which is a sign of overcorrection. L) sva shows little change from the raw data. The batch variable still shows separation with the GE_PHILLIPS more towards the bottom right and SIEMENS more towards the top left. **kBET analysis:** M) Raw data has a significant difference between the observed and expected rejection rate (p=0) indicating that a batch effect is present. N) ComBat corrected data does not have a significant difference between the observed and expected rejection rate (p=0.05), indicating that the batch effect has been corrected. O) limma/voom corrected data does not have a significant difference between the observed and expected rejection rate (p=0.55), indicating that the batch effect is no longer present. P) sva corrected data has a significant difference between the observed and expected rejection rate (p=0) indicating that a batch effect is still present.

Furthermore, in the kBET analysis, the observed rejection rate (mean = 0.624) is significantly (p = 0) greater than the expected rejection rate (mean = 0.008) (Figure 5M). Therefore, there is likely a batch effect that we should correct.

##### Batch Correct

Since the ADNI dataset contains nondiscrete data, the AIC GoF will only calculate lognormal and voom scores. The lowest AIC score is for voom (−495.6981), indicating that the data may fit a normal rather than a lognormal distribution (AIC score = Inf, indicating that no single gene had the lowest AIC value for a lognormal distribution; See Supplemental Figure 6). Additionally, the negative binomial GoF indicates that a negative binomial distribution may be appropriate, as 0% of the features are significant. Therefore, we will consider utilizing ComBat, voom/limma, or sva as correction methods. Since this is not RNA-Seq data, ComBat-Seq would not be a good method. We apply a log normalization prior to ComBat and sva correction and a voom normalization prior to limma correction.

##### Analyze Batch Correction Methods

In the variation analysis, ComBat shows a reduction in the batch effect, with “Manufacturer” (mean = 0.04) explaining significantly less variation (p < 0.001) than “Diagnosis” (mean = 11.8) (Figure 5B). voom/limma shows overcorrection since the explained variation is not attributed to either the batch (mean = 0) or biological (mean = 1.29) variable (Figure 5C) and sva does not seem to show any difference when compared to the raw analysis (batch mean = 7.4, biological mean = 11.1, p = 0.16) (Figure 5D). In terms of λ, the raw λ was 0.74 and both ComBat (λ = -3.54) and limma/voom (λ = -56.31) provide better rates. However, limma/voom has a drastically smaller λ, another sign of overcorrection. The sva λ is 1.12, worse than the raw λ, indicating that sva is likely not a good option for correction. ComBat displays homogenous mixing of the “Manufacturer” batch variable in both the PCA plot (Figure 5F) and the UMAP (Figure 5J). voom/limma also shows homogenous mixing of the batch variable in both the PCA plot (Figure 5G) and the UMAP (Figure 5K). Again, sva still shows clustering by the batch variable in both the PCA (Figure 5H) and the UMAP (Figure 5L). Finally, the kBET analysis for ComBat (Figure 5N) shows a large decrease in the mean observed rate (mean = 0.114), making it statistically comparable (p = 0.05) to the expected rate (mean = 0.035). voom/limma follows the same pattern, with the observed rate (mean = 0.038) closer to the expected rate (mean = 0.041; Figure 5O), making the results not significantly different (p = 0.55) and indicating that the batch effect has been corrected. However, for sva, the observed rate (mean = 0.644) is still significantly (p = 0) greater than the observed rate (mean = 0.018; Figure 5P).

These results indicate that ComBat is likely the best choice for batch correction as it reduces the batch effect while maintaining the biological variable. Voom/limma could also be a good choice, but there is a risk of eliminating the variance due to the biological variable.

## 4 Discussion

As our TB and ADNI examples indicate, the need for batch correction spans a wide range of data types, not limited to -omics data. BatchQC allows users to determine the need for batch effect using a variety of visualization and evaluation methods, including more traditional methods such as dimension reduction and novel methods, including our λ statistic and goodness-of-fit. After deciding that a batch correction is needed, users can normalize and correct their data within BatchQC and then analyze which correction method is best suited for their data. All normalized and corrected datasets can be exported as a SummarizedExperiment object and used directly with many downstream tools. These steps can be performed through the shiny app, or with the functions provided in the BatchQC package.

In general, our recommendation is that unbalanced designs with an experimental batch effect usually need to be corrected, regardless of the extent of the unbalanced design (See Figures 1B and 1C). However, balanced designs only need to be corrected when the variance explained by the batch exceeds a λ value of -7 (see Figure 1A). The λ value can also be used to compare raw, uncorrected datasets with different correction methods to determine which method provides the best λvalue.

kBET^13^ is a published method that has been previously used in existing batch effect comparisons^32^, but to our knowledge, has not been implemented in other batch correction tools. After applying various batch effect correction methods, researchers can use kBET to quantitatively compare the effectiveness of these methods. By calculating rejection rates before and after correction, or by comparing rejection rates across different correction approaches, kBET provides an objective benchmark for determining which method most successfully removes batch effects while preserving biological variation. This quantitative comparison is particularly valuable when choosing between multiple correction strategies or optimizing correction parameters. Within our ADNI dataset, kBET confirmed that ComBat was the best correction method as indicated by multiple visualization methods. However, like other methods, it is best utilized in conjunction with other methods (as seen by the contradictory results in the TB example data).

Furthermore, the AIC comparison can be used to assess the distribution that best fits a dataset, informing general downstream analysis as well as selecting an appropriate batch effect correction. We calculate an AIC score for the negative binomial, log-normal, and normal distributions for each feature in a dataset. We recommend using the distribution that has the lowest AIC score (𝛽i) as the appropriate distribution for the dataset as a whole as this metric aids in preventing excessive impact of a few major outlier genes. The additional insight that the AIC comparison provides regarding the distribution of the users’ dataset can enable users to more appropriately select the best batch correction method and other downstream tools. This feature is widely applicable outside of batch effect correction analyses, such as for determining the appropriate method for differential expression.^31,33,34^

When specifically using tools that require a negative binomial distribution, our negative binomial GoF can assess that the dataset meets the model distribution assumption for edgeR^15^ (which uses a negative binomial distribution). Our model-based GoF method is more suitable for smaller datasets (less than or equal to 20 samples), where permutation-based methods may lack statistical power^31^. The model-free method proposed by Li et al. is recommended for larger datasets (20 or more samples), where the effectiveness of permutation tests improves with increased sample size^31^. By utilizing the method appropriate for the sample size, researchers can assess whether the negative binomial model assumptions hold and whether algorithms relying on the negative binomial distribution produce valid results (for example, ComBat-Seq also uses a similar model formulation^23^).

Since no single method can properly assess all cases of batch effect as seen in our TB and ADNI data examples, BatchQC includes multiple methods, including both visualization and quantitative methods, to fit a wide range of cases. We recommend that researchers thoroughly explore their data before determining the most suitable batch correction method. BatchQC includes multiple tools to aid researchers in this process, including adding ComBat-Seq^23^, Harman^11^, limma^12^, and sva-seq^25^ as batch correction strategies, upgraded PCA plots, and the addition of UMAPs, GoF tools, and the λ statistic. Ultimately, researchers will need to use their best judgment in applying correction techniques and using downstream tools.

Additionally, the BatchQC R program was written to allow a flexible design centered around object-oriented programming. This enables all functions utilized in the Shiny R app to be accessed and used from the command line, independent of app utilization. This creates flexibility, such as using BatchQC features in an analysis pipeline. This also provides an effortless means to document the functions and parameters used in an analysis, allowing for more streamlined reproducibility efforts. Further, the object-oriented nature of BatchQC allows users to dynamically upload a dataset, enabling them to analyze multiple datasets within a single session of the Shiny app. Finally, once all analyses for a dataset are complete, the user can download a SummarizedExperiment object with all related assays created through normalization or batch correction steps. This SE object can then be utilized as input to other downstream tools.

We acknowledge that BatchQC has limitations, including not having all possible batch correction tools currently available as correction options (Table 1), our new AIC GoF has limited functionality for non-discrete datasets, and batch correction best practices continue to expand and may not be reflected in the tools available in BatchQC. However, BatchQC is a powerful tool for exploring and analyzing datasets when applying batch correction.

As we look toward the future of batch correction, we anticipate expanding the available batch correction tools in BatchQC, including those specific to datasets beyond RNA-Seq, such as single-cell and methylation data (e.g., ComBat-met^35^). Furthermore, we would like to create workflows tailored to each of these data types to streamline the user experience.

## 5 Data Availability

All simulation data and method data, code for methods, results, and figures are available at github.com/jessmcc22/BatchQCv2_Manuscript except for the data needed for the negative binomial simulation, which is available at https://doi.org/10.5281/zenodo.17429046.

## 6 Code Availability

BatchQC is an open-source software; the most up-to-date version is available for download through Bioconductor. All software code, including the development version, is also available on GitHub at github.com/wejlab/BatchQC. All software was written in R version 4.5.0. All code for methods, results, and figures in this paper is available at the Manuscript GitHub (github.com/jessmcc22/BatchQCv2_Manuscript).

## 7 Acknowledgments

We would like to acknowledge support received from R01 grant GM127430 and R01 MH123550.

The results published here are in part based upon data generated by the TCGA Research Network: https://www.cancer.gov/tcga. The Genotype-Tissue Expression (GTEx) Project was supported by the Common Fund of the Office of the Director of the National Institutes of Health, and by NCI, NHGRI, NHLBI, NIDA, NIMH, and NINDS. The data used for the analyses described in this manuscript were obtained from the GTEx Portal in 2021 in conjunction with Li et al.’s previous paper^31^.

Additional data collection and sharing for this project was funded by the Alzheimer’s Disease Neuroimaging Initiative (ADNI) (National Institutes of Health Grant U01 AG024904) and DOD ADNI (Department of Defense award number W81XWH-12-2-0012). ADNI is funded by the National Institute on Aging, the National Institute of Biomedical Imaging and Bioengineering, and through generous contributions from the following: AbbVie, Alzheimer’s Association; Alzheimer’s Drug Discovery Foundation; Araclon Biotech; BioClinica, Inc.; Biogen; Bristol-Myers Squibb Company; CereSpir, Inc.; Cogstate; Eisai Inc.; Elan Pharmaceuticals, Inc.; Eli Lilly and Company; EuroImmun; F. Hoffmann-La Roche Ltd and its affiliated company Genentech, Inc.; Fujirebio; GE Healthcare; IXICO Ltd.; Janssen Alzheimer Immunotherapy Research & Development, LLC.; Johnson & Johnson Pharmaceutical Research & Development LLC.; Lumosity; Lundbeck; Merck & Co., Inc.; Meso Scale Diagnostics, LLC.; NeuroRx Research; Neurotrack Technologies; Novartis Pharmaceuticals Corporation; Pfizer Inc.; Piramal Imaging; Servier; Takeda Pharmaceutical Company; and Transition Therapeutics. The Canadian Institutes of Health Research is providing funds to support ADNI clinical sites in Canada. Private sector contributions are facilitated by the Foundation for the National Institutes of Health (www.fnih.org). The grantee organization is the Northern California Institute for Research and Education, and the study is coordinated by the Alzheimer’s Therapeutic Research Institute at the University of Southern California. ADNI data are disseminated by the Laboratory for Neuro Imaging at the University of Southern California.

## 8 Author Contributions

JKA implemented and oversaw the bulk of the restructuring and new analysis tools introduced in BatchQC, completed TB and ADNI data analysis with BatchQC, compiled all methods, and wrote this paper and all supporting documents. JZ created all methods associated with the AIC comparison and lambda statistic. TL oversaw the creation of the AIC comparison and lambda statistic, provided written portions to support these areas of the paper, and collaborated extensively with JKA and WEJ to create and implement these methods. XG created the negative binomial GoF and provided written methods for these sections. HF implemented kBET in BatchQC, wrote sections of the paper containing to kBET, and reviewed the manuscript code. YL implemented many of the new tools available, including the AIC comparison, sva and sva-seq batch correction, and Kruskal-Wallis and ANOVA differential expression. MS, RC, ZL, and EH developed the initial new architecture for BatchQC, which enabled object-oriented programming. EH additionally implemented the circular dendrograms and volcano plot. SSJ updated the documentation, added examples to functions that previously lacked them, and enhanced the visualization of heatmaps and dendrograms. SL implemented the Harman batch correction method. RTS provided the data for the ADNI analysis and collaborated on using BatchQC tools on imaging data. WEJ oversaw the creation and implementation of all statistical tools and collaborated in depth with ZG, RTS, and TL. All authors reviewed this manuscript.

## References

1 Manimaran, S. et al. BatchQC: interactive software for evaluating sample and batch effects in genomic data. Bioinformatics 32, 3836–3838 (2016). 10.1093/bioinformatics/btw538

2 Davis, J. T. et al. BatchFLEX: feature-level equalization of X-batch. Bioinformatics 40 (2024). 10.1093/bioinformatics/btae587

3 Zhu, T. et al. BatchServer: A Web Server for Batch Effect Evaluation, Visualization, and Correction. Journal of Proteome Research 20 (2020). 10.1021/acs.jproteome.0c00488

4 Gebreyesus, G. et al. Combining multi-population datasets for joint genome-wide association and metaanalyses: The case of bovine milk fat composition traits. Journal of Dairy Science 102 (2019). 10.3168/jds.2019-16676

5 Hamid, J. S. et al. Data Integration in Genetics and Genomics: Methods and Challenges. Human Genomics and Proteomics : HGP 2009 (2009). 10.4061/2009/869093

6 Leek, J. T. et al. Tackling the widespread and critical impact of batch effects in high-throughput data. Nat Rev Genet 11, 733–739 (2010). 10.1038/nrg2825

7 Kucukural, A. et al. DEBrowser: interactive differential expression analysis and visualization tool for count data. BMC Genomics 2019 20:1 20 (2019). 10.1186/s12864-018-5362-x

8 Johnson, W. E., Li, C. & Rabinovic, A. Adjusting batch effects in microarray expression data using empirical Bayes methods. Biostatistics 8, 118–127 (2007). 10.1093/biostatistics/kxj037

9 Yu, Y., Mai, Y., Zheng, Y. & Shi, L. Assessing and mitigating batch effects in large-scale omics studies. Genome Biology 25 (2024). 10.1186/s13059-024-03401-9

10 Chang W, C. J., Allaire J, Sievert C, Schloerke B, Aden-Buie G, Xie Y, Allen J, McPherson J, Dipert A, Borges B. shiny: Web Application Framework for R. (2025).

11 Oytam, Y. et al. Risk-conscious correction of batch effects: maximising information extraction from highthroughput genomic datasets. BMC Bioinformatics 2016 17:1 17 (2016-09-01). 10.1186/s12859-016-1212-5

12 Ritchie, M. E. et al. limma powers differential expression analyses for RNA-sequencing and microarray studies. Nucleic Acids Research 43 (2015). 10.1093/nar/gkv007

13 Büttner, M. et al. A test metric for assessing single-cell RNA-seq batch correction. Nature Methods 16, 43–49 (2019). 10.1038/s41592-018-0254-1

14 Law, C. W. et al. voom: precision weights unlock linear model analysis tools for RNA-seq read counts. Genome Biology 2014 15:2 15 (2014). 10.1186/gb-2014-15-2-r29

15 Chen, Y., Chen, L., Lun, Aaron T. L., Baldoni, Pedro L. & Smyth, Gordon K. edgeR v4: powerful differential analysis of sequencing data with expanded functionality and improved support for small counts and larger datasets. Nucleic Acids Research 53 (2025). 10.1093/nar/gkaf018

16 VanValkenburg, A. et al. Malnutrition leads to increased inflammation and expression of tuberculosis risk signatures in recently exposed household contacts of pulmonary tuberculosis PubMed. Frontiers in immunology 13 (2022). 10.3389/fimmu.2022.1011166

17 NJ, T., et al. Longitudinal Mapping of Cortical Thickness Measurements: An Alzheimer’s Disease Neuroimaging Initiative-Based Evaluation Study PubMed. Journal of Alzheimer’s disease : JAD 71 (2019). 10.3233/JAD-190283

18 Hu, F. et al. DeepComBat: A Statistically Motivated, Hyperparameter-Robust, Deep Learning Approach to Harmonization of Neuroimaging Data. bioRxiv (2023). 10.1101/2023.04.24.537396

19 Morgan M, O. V., Hester J, Pagès H. SummarizedExperiment: A container (S4 class) for matrix-like assays. (2025). doi:10.18129/B9.bioc.SummarizedExperiment

20 Soneson, C. et al. Eleven quick tips for writing a Bioconductor package. PLOS Computational Biology 21 (Mar 19, 2025). 10.1371/journal.pcbi.1012856

21 Love, M. I. et al. Moderated estimation of fold change and dispersion for RNA-seq data with DESeq2. Genome Biology 2014 15:12 15 (2014). 10.1186/s13059-014-0550-8

22 Ramos, M. et al. Software for the Integration of Multiomics Experiments in Bioconductor. Cancer Research 77 (2017/11/01). 10.1158/0008-5472.CAN-17-0344

23 Zhang, Y., Parmigiani, G. & Johnson, W. E. ComBat-seq: batch effect adjustment for RNA-seq count data. NAR Genom Bioinform 2, lqaa078 (2020). 10.1093/nargab/lqaa078

24 Leek, J. T., Johnson, W. E., Parker, H. S., Jaffe, A. E. & Storey, J. D. The sva package for removing batch effects and other unwanted variation in high-throughput experiments. Bioinformatics 28 (2012 Jan 17). 10.1093/bioinformatics/bts034

25 Leek, J. T. svaseq: removing batch effects and other unwanted noise from sequencing data. Nucleic Acids Research 42 (2014/12/01). 10.1093/nar/gku864

26 Wickham, H. ggplot2: Elegant Graphics for Data Analysis. (2016).

27 R, K. pheatmap: Pretty Heatmaps. (2025).

28 Vries, A. d. & Ripley, B. D. Create Dendrograms and Tree Diagrams Using ’ggplot2’. (2025).

29 Team, R. C. R: A Language and Environment for Statistical Computing. (2025).

30 Konopka, T. umap: Uniform Manifold Approximation and Projection. (2023). 10.32614/CRAN.package.umap

31 Li, Y. et al. Exaggerated false positives by popular differential expression methods when analyzing human population samples. Genome Biology 2022 23:1 23 (2022). 10.1186/s13059-022-02648-4

32 Tran, H. T. N. et al. A benchmark of batch-effect correction methods for single-cell RNA sequencing data. Genome Biology 2020 21:1 21 (2020-01-16). 10.1186/s13059-019-1850-9

33 Lin, Y. et al. Comparison of normalization and differential expression analyses using RNA-Seq data from 726 individual Drosophila melanogaster. BMC Genomics 2016 17:1 17 (2016). 10.1186/s12864-015-2353-z

34 Hawinkel, S., Rayner, J. C. W., Bijnens, L. & Thas, O. Sequence count data are poorly fit by the negative binomial distribution. PLOS ONE 15 (Apr 30, 2020). 10.1371/journal.pone.0224909

35 Wang, J. ComBat-met: adjusting batch effects in DNA methylation data. NAR Genomics and Bioinformatics 7 (2025/03/29). 10.1093/nargab/lqaf062

