## Supplemental Figures for "Reproducible Tools and Enhanced Computational Workflows for Batch Effect Evaluation of High-Throughput Data Using BatchQC"

### BatchQC Supplemental Figures

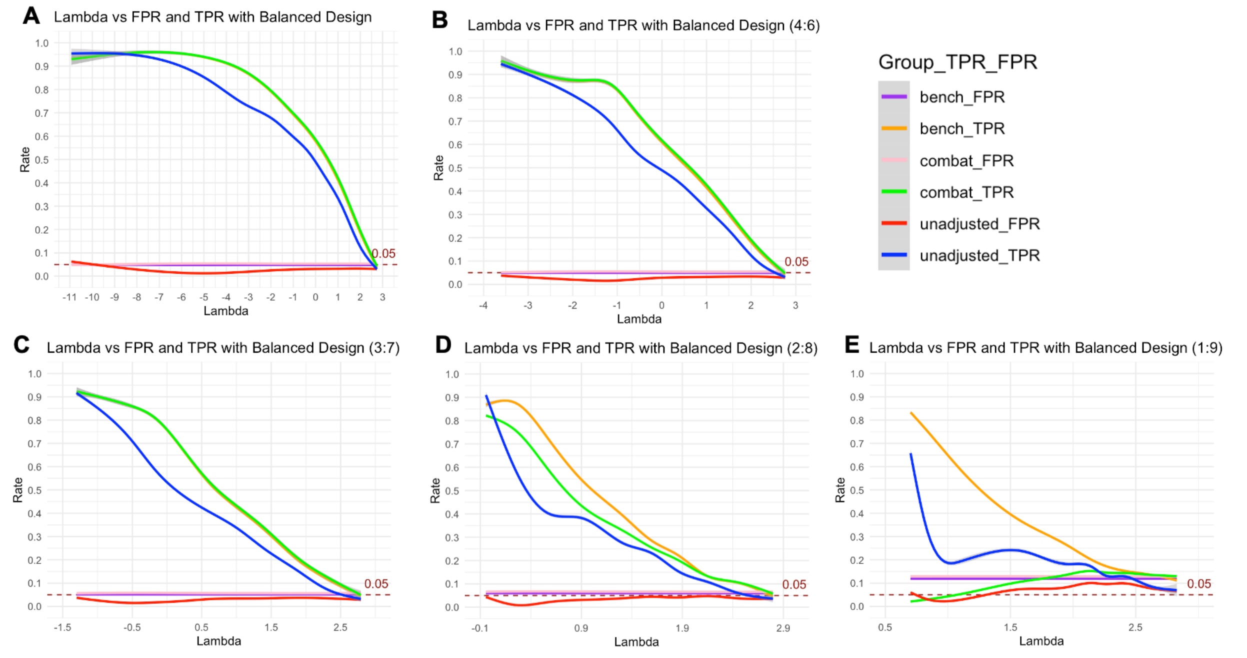


**Supplemental Figure 1: False positive and true positive rates based on balanced designs in different ratios.** A) Balanced design in a 5:5 ratio. B) Balanced design in a 4:6 ratio between batches. C) Balanced design in a 3:7 ratio between batches. D) Balanced design in a 2:8 ratio between batched. E) Balanced design in a 1:9 ratio between batches. As the ratio between batches becomes more unbaanced, correction for batch helps maintain a statistically valid FPR (E). As lambda gets larger, correcting for batch maintains a higher TPR.

A

| AIC metric | NB | Lognormal | Voom |
|---|---|---|---|
| total AIC | 1561349.9 | 1940302.9 | 867822.3 |
| min AIC | 658 | 1 | 128 |
| AIC score | 2372.872 | 1940302.922 | 6779.862 |
| Median AIC | 312.9978 | 2494.1958 | 1062.9028 |

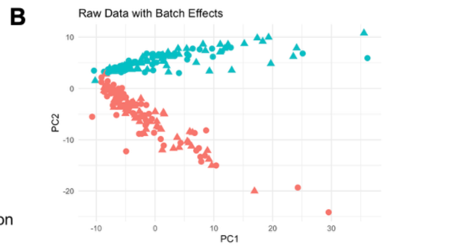


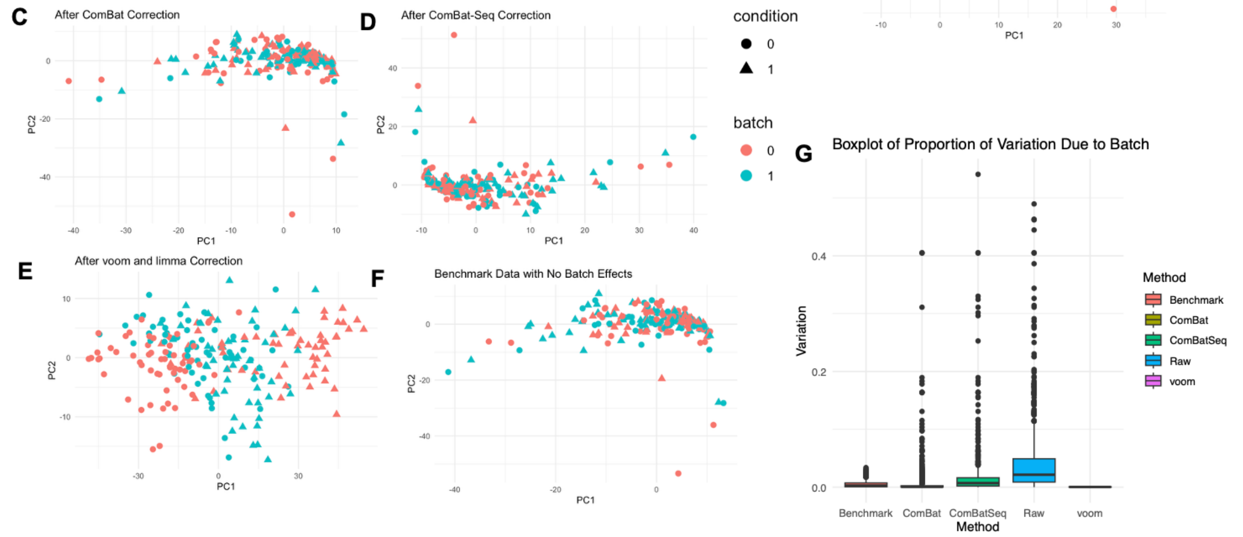


**Supplemental Figure 2: AIC model fit on negative binomial simulated data**. The AIC score correctly predicts that the simulated dataset best fits a negative binomial model (A) since the lowest AIC score is for the negative binomial model. PCA plots (B-F) plot the condition variable (red or blue) and the batch (circle or triangle) to show groupings prior to batch correction on the simulated data(B), after ComBat correction (C), after ComBat-Seq correction (D), after Voom and limma correction (E), and the benchmark data with no batch effect and no correction (F). We see that all three methods reduce the grouping based on batch, however, note that both ComBat and voom/limma overcorrect the data as seen in the boxplot (G). Both methods have less variation than the benchmark data which has no batch effect. Whereas ComBat-Seq reduces the overall variation due to batch effect without overcorrection. This supports the suggestion that the negative binomial is the best fit for the data and thus, ComBat-Seq would be an appropriate correction method to utilize.

A

| AIC metric | NB | Lognormal | Voom |
|---|---|---|---|
| total AIC | 1318607.8 | 943333.9 | 576205.2 |
| min AIC | 1 | 496 | 290 |
| AIC score | 1318607.8 | 1901.883 | 1986.915 |
| Median AIC | 1668.2668 | 633.6008 | 722.1260 |

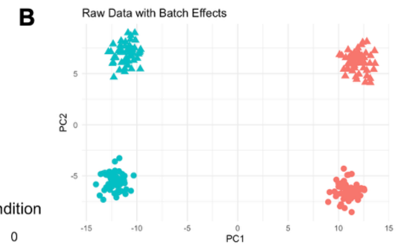


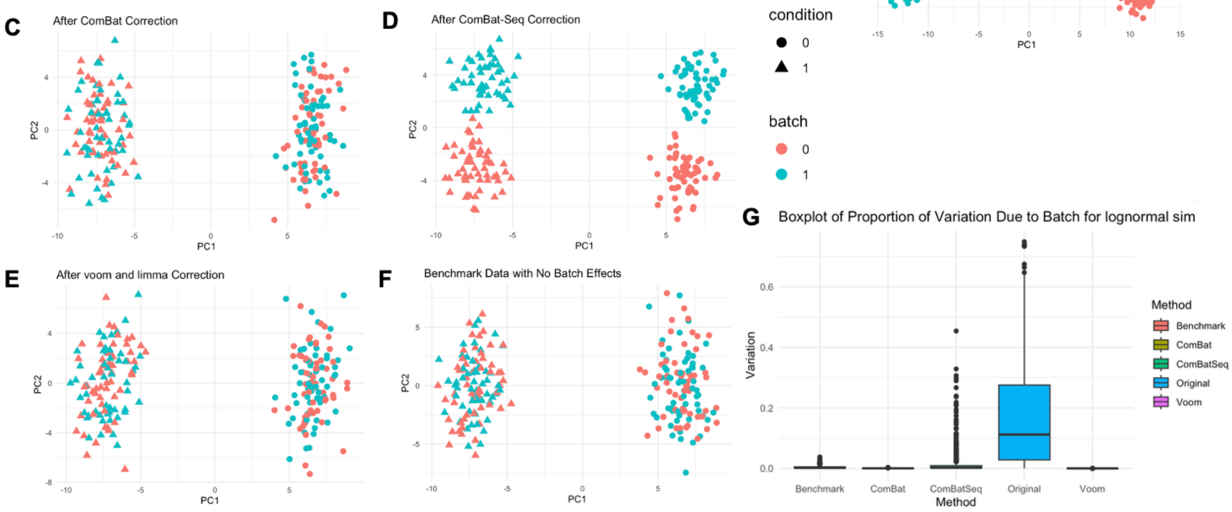


**Supplemental Figure 3: AIC model fit on lognormal simulated data**. The AIC tool correctly predicts that the simulated dataset best fits lognormal model (A) since the lowest AIC score is for the lognormal model. PCA plots (B-F) plot the condition variable (red or blue) and the batch (circle or triangle) to show groupings prior to batch correction on the simulated data (B), after ComBat correction (C), after ComBat-Seq correction (D), after Voom and limma correction (E), and the benchmark data with no batch effect and no correction (F). We see that all three methods reduce the grouping based on batch; however, Combat-Seq (D) still has distinct batch groupings. ComBAt (C) and Voom and limma (E) provide the best mixing of batches. The boxplot indicates that each method reduces the variation to be similar to the benchmark data (G).

A

| AIC metric | NB | Lognormal | Voom |
|---|---|---|---|
| total AIC | 1262320.4 | 1398355.9 | 540453.8 |
| min AIC | 198 | 312 | 277 |
| AIC score | 6375.356 | 4481.910 | 1951.097 |
| Median AIC | 1460.0746 | 2344.2438 | 687.6223 |

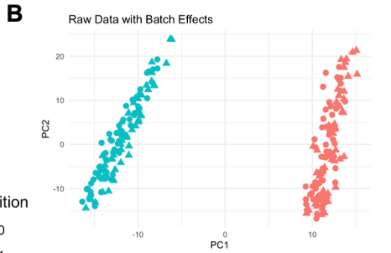


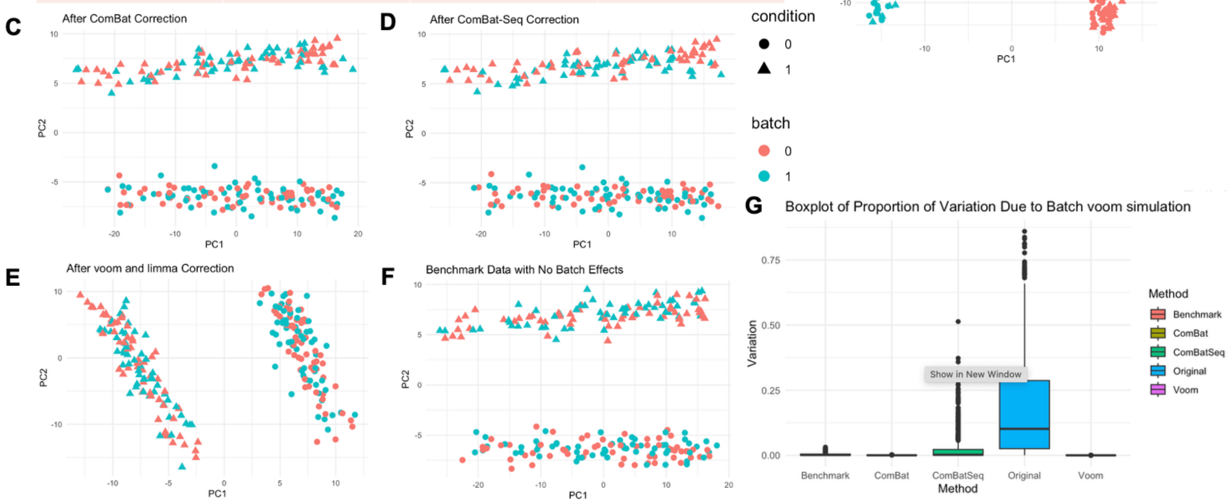


**Supplemental Figure 4: AIC model fit on voom simulated data**. The AIC tool correctly predicts that the simulated dataset best fits a normal distribution (A) since the lowest AIC score is for the voom model. PCA plots (B-F) plot the condition variable (red or blue) and the batch (circle or triangle) to show groupings prior to batch correction on the simulated data (B), after ComBat correction (C), after ComBat-Seq correction (D), after Voom and limma correction (E), and the benchmark data with no batch effect and no correction (F). We see that all three methods reduce the grouping based on batch. Additionally, the boxplot indicates that each method reduces the variation to be similar to the benchmark data (G).

A

| AIC metric | NB | Lognormal | Voom |
|---|---|---|---|
| Total AIC | 18608300.213 | 4683915.6234 | 3174420.1261 |
| Freq. of min AIC | 1 | 14109 | 3247 |
| AIC score | 18608300.213 | 331.9807 | 977.6471 |
| Median AIC | 1099.963 | 128.2294 | 177.1217 |

B

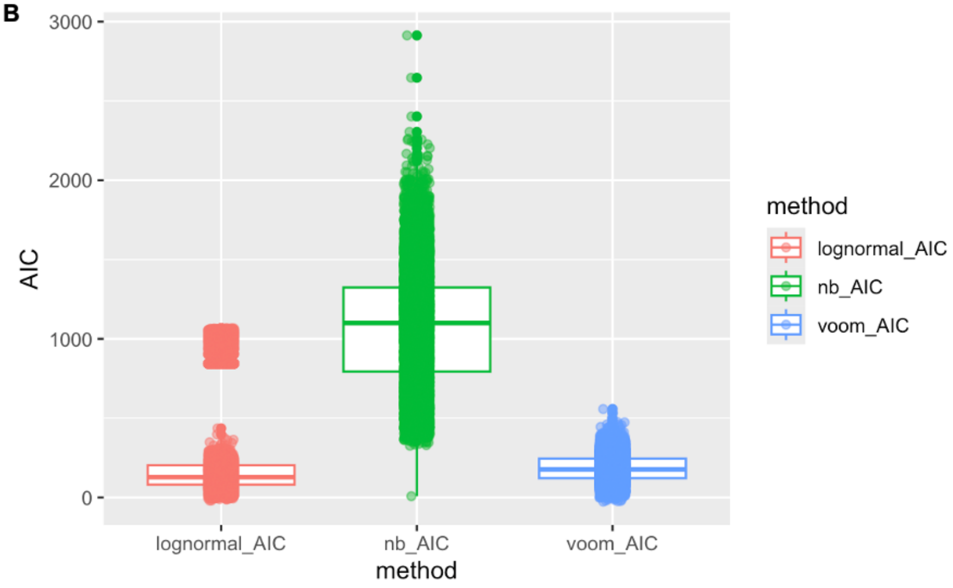


**Supplemental Figure 5: TB AIC Results.** A) The lowest AIC score if for lognormal (331.9807). B) We see similar distributions for lognormal and voom with a number of outliers for lognormal. However, more genes have a lower AIC score in the lognormal model (A, Freq of min AIC = 14109) compared to voom (A, Freq of min AIC = 3247), indicating that lognormal is likely a good fit.

A

| AIC metric | Lognormal | Voom |
|---|---|---|
| Total AIC | 1116.01582 | -3965.5847 |
| Freq. of min AIC | 0 | 8 |
| AIC score | Inf | -495.6981 |
| Median AIC | 39.20847 | -636.9979 |

B

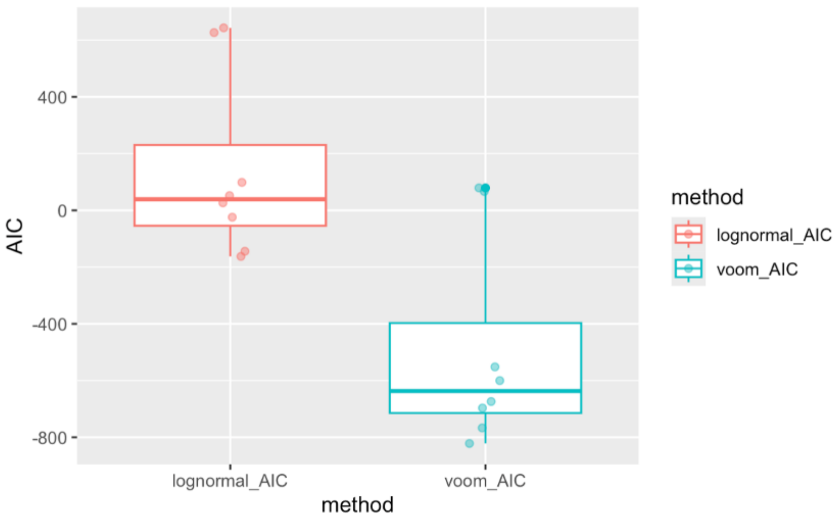


**Supplemental Figure 6: ADNI AIC Results.** A) The lowest AIC score if for voom (-495.6981). B) The distribution of AIC scores is lower for voom than for lognormal, other than one outlier. This indicates that a voom normal distribution is likely a better fit that lognormal.
